# mdBIRCH for Fast, Scalable, Online Clustering of Molecular Dynamics Trajectories

**DOI:** 10.1101/2025.11.05.686879

**Authors:** Jherome Brylle Woody Santos, Lexin Chen, Ramón Alain Miranda-Quintana

## Abstract

We present mdBIRCH, an online clustering method that adapts the BIRCH CF-tree to molecular dynamics (MD) data by applying a merge test calibrated directly to RMSD. Each arriving frame is routed to the nearest leaf microcluster and merged only if the post-merge centroid-based spread, computed from the cluster feature (CF) summaries, remains within a user-supplied threshold *τ*. This enables incremental, memory-bounded operation without constructing pairwise distance matrices, with a physically interpretable parameter controlling structural granularity. We evaluate mdBIRCH on *β*-heptapeptide and HP35 systems and propose two practical strategies to make the threshold selection easier: (a) RMSD-anchored runs that use controlled structural edits to define interpretable operating points and (b) a blind sweep that tracks how cluster counts, occupancies, and coverage evolve with the threshold. Across both systems, increasing this threshold produces predictable consolidation into fewer, higher-occupancy states, and broader RMSD-to-centroid distributions. We further analyze the effect of the CF-tree capacity through the branching factor, quantify sensitivity to data ordering, and compare dominant-state representatives against batch clustering workflows to contextualize the resulting partitions. Finally, because decisions rely only on cluster summaries, mdBIRCH scales near-linearly with the number of frames on standard CPU hardware, offering a practical combination of speed and interpretability for large-scale trajectory analysis.

## 1. INTRODUCTION

Clustering remains one of the most practical ways to make sense of large molecular dynamics (MD) trajectories.^1–3^ By grouping structurally similar frames, one can pinpoint the dominant states or pick representatives for visualization or refinement.^3,4^ Despite its usefulness, scalability is one of its persistent limitations. Many classical approaches rely on pairwise distance information, explicitly through distance matrices or implicitly through repeated neighborhood queries, so their time and memory costs can grow rapidly as trajectories reach hundreds of thousands to millions of frames. In practical workflows, this often forces downsampling (sieving) or other approximations that reduce the number of frames. While useful for speed, these strategies can miss rare but meaningful conformations.

In our recent work on Hierarchical Extended Linkage Method (HELM), we showed that it is possible to avoid the full quadratic O(*N*^*2*^) cost of pairwise distance matrices while still producing high-quality hierarchical clustering outputs.^5^ In HELM, we first group the frames into microclusters and then construct the hierarchy over all subclusters using *n*-ary calculations. These *n*-ary similarity functions^6–13^ let us evaluate linkage relationships between subclusters directly, without enumerating all pairwise distances, while keeping every frame assigned through its subcluster membership. The result is hierarchy-quality clustering that remains manageable, even at the scale of million-frame trajectories.

There are, however, common scenarios where a different tool would be more valuable than a batch pipeline. First, trajectories are produced incrementally: simulations extend over time, new trajectories may be appended, new ensembles may be launched, or new segments may arrive from adaptive sampling.^14–17^ In these settings, it is convenient to be able to update clustering results without rebuilding a global model from scratch each time new frames are generated. Second, it is appealing in practice to tune clustering with only one interpretable parameter rather than multiple coupled hyperparameters such as the number of clusters, minimum points, neighborhood radius, linkage rules, and the like. Third, users often want a simple, bounded notion of within-cluster spread.^18–20^ In many analyses, it is useful to be able to ensure that clusters do not become arbitrarily broad as more frames are assigned (i.e., to keep the cluster’s centroid-based average measure of spread within a specified bound), especially when clusters are used to choose representatives or to define states for downstream modeling.

These needs motivate mdBIRCH. We adapt the classic BIRCH (Balanced Iterative Reducing and Clustering using Hierarchies)^21^ framework to molecular dynamics trajectories. BIRCH was designed for large datasets and uses a compact summary of each cluster, called a cluster feature (CF), to support fast insertion of points into a tree structure (CF-tree) without needing all prior points. In mdBIRCH we retain this streaming architecture but replace the usual geometric spread criterion with one calibrated directly to widely used RMSD (root mean square deviation). In essence, mdBIRCH tests each proposed assignment by computing the candidate cluster’s spread after adding the new frame (post-merge) and accepts the assignment if and only if the resulting spread remains within a user-specified threshold *τ*.

The result is a method that offers: (a) incremental operation: we can update clusters as new frames arrive, rather than requiring a complete trajectory in advance; (b) a primary interpretable parameter (in RMSD units) that we can use to represent cluster tightness; and (c) scalability for large trajectories: because clustering decisions are made from CF summaries the method is practical for long MD runs. In this work, we derive the RMSD-calibrated acceptance rule, describe practical strategies for selecting *τ*, and benchmark mdBIRCH to evaluate clustering behavior and computational efficiency.

## 2. THEORY

### 2.1 Frame representation and CF summaries

We assume that the trajectory frames have been aligned to a reference using the same atom selection used for RMSD calculations. After alignment, each frame can be represented as a vector *x* in ℝ^*D*^ where *D* is the number of coordinates/features (e.g., *D* = 3 × number of selected atoms).

Rather than storing all frames assigned to a cluster, mdBIRCH summarizes each cluster using a cluster feature (CF), which stores sufficient statistics to update the centroid and compute its centroid-based measure of spread. Following the BIRCH framework, we define the CF for cluster *j* as Eq. (1):

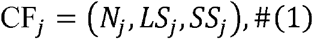

where *N*_*j*_ is the number of frames in the cluster, *LS*_*j*_ is the component-wise linear sum of the frame vectors given by Eq. (2), *SS*_*j*_ and is the scalar sum of the squared norms defined by Eq.(3):

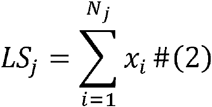

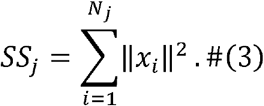

From these summaries, the cluster centroid is computed using Eq. (4):

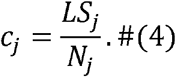

This is the key computational idea: when a new frame arrives, we do not recompute quantities over all members of the cluster. Instead, we update the CF in *O*(*D*)without revisiting stored cluster members.

### 2.2 Centroid-based spread from CFs

To decide whether an incoming frame *x*_*i*_ should be merged into a candidate cluster, mdBIRCH uses the mean squared radius about the cluster centroid as shown in Eq. (5):

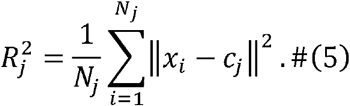

Using the CF summaries, this quantity can be computed without revisiting stored frames using Eq. (6):

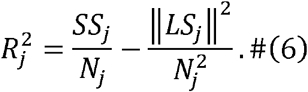

Intuitively, the CF is a compact “memory” of a cluster that supports fast updates. When a new frame arrives, mdBIRCH can test whether adding it would keep the cluster sufficiently tight using only cheap operations on the CF rather than distances between all pairs of frames.

For interpretability across atom selections of different dimensionality, we report the per-coordinate centroid variance shown in Eq. (7):

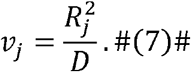

When inserting a new frame *x* into cluster *j*, the updated CF is obtained by Eq. (8):

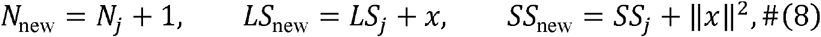

and the corresponding post-merge spread is computed from Eq. (6) using (*N*_new_, *LS*_new_, *SS*_new_ ).

### 2.3 RMSD-calibrated merge criterion

To make the threshold parameter interpretable in familiar RMSD units, we relate the centroid-based spread to the RMSD between two frames. Throughout, we define RMSD from reference-aligned coordinate vectors according to Eq. (9):

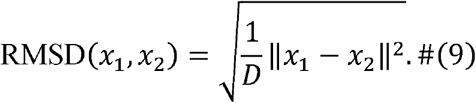

We then consider a two-frame cluster nucleated by two aligned frames *x*_1_ and *x*_2_, whose centroid *c* = (*x*_1_ + *x*_2_ ) /2. For this cluster, the spread is calculated as Eq. (10):

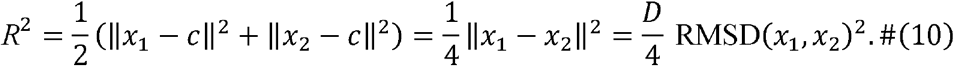

This yields a direct relationship between the centroid-based spread and an RMSD-scale threshold.

Accordingly, for a tentative insertion into an existing cluster, mdBIRCH forms the hypothetical post-merge CF and recomputes the post-merge centroid spread from it using Eq. (11):

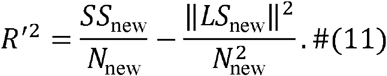

We therefore accept the merge whenever the post-merge mean squared radius satisfies Eq. (12):

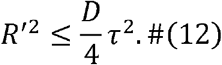

Here, *τ* is the RMSD-calibrated threshold: in the two-frame case, Eq. (12) is exactly equivalent to requiring RMSD((*x*_1_, *x*_2_ ) ≤ *τ*. For larger clusters, the same rule bounds the centroid-based average spread under post-merge updates. Full derivation can be found in Supporting Information.

### 2.4. Decision granularity and interpretation

Operationally, as illustrated in Fig. 1, each incoming frame is routed down the CF-tree to a candidate leaf microcluster (nearest centroid). The algorithm then evaluates the hypothetical merged CF (i.e., the CF after adding the new frame), recomputes the post-merge spread, and applies the acceptance rule. If accepted, the microcluster CF is updated; otherwise, a new microcluster is created.

**Figure 1.**
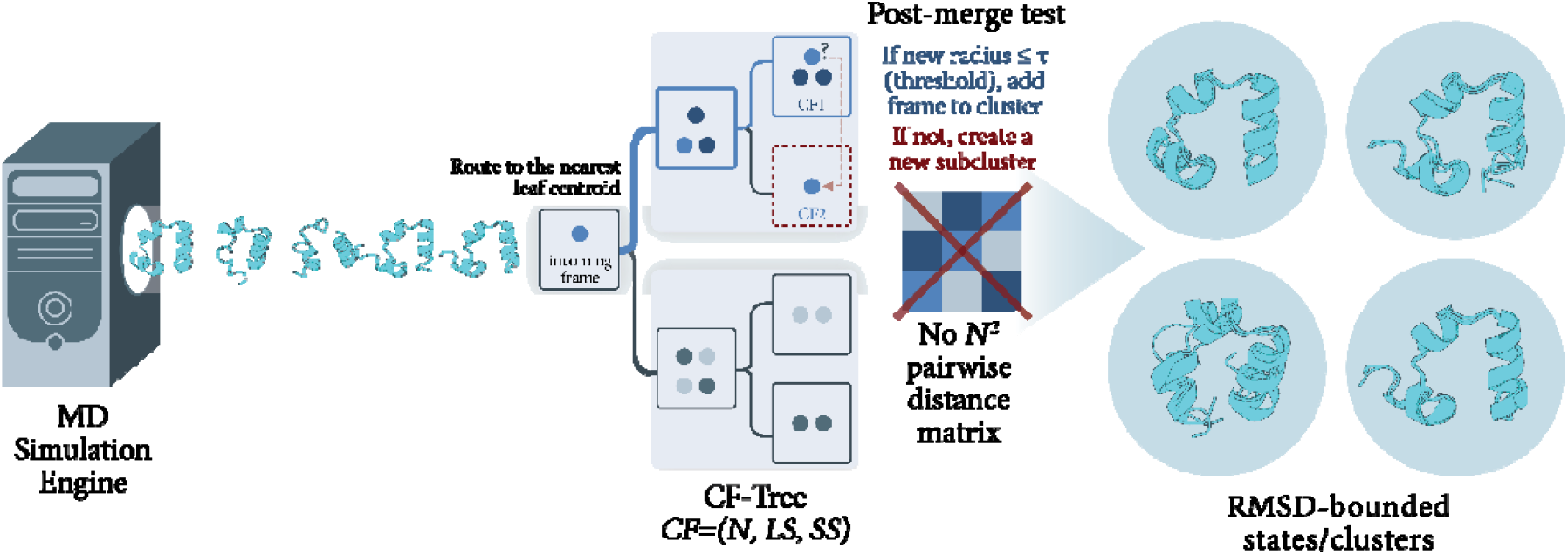
Schematic of mdBIRCH streaming insertion. Frames are processed sequentially and routed to the nearest leaf microcluster in a CF-tree. A post-merge test computes the merged centroid-based spread and accepts the insertion if the new measure is within the user-specified threshold; otherwise, a new microcluster is created.

The CF-tree is controlled by the branching factor (BF), which sets the maximum number of CF entries stored in an internal node and the maximum number of microcluster entries stored in a leaf. When an insertion would cause a node to exceed its capacity, the node is split following the standard BIRCH procedure. This splitting keeps the tree shallow, supporting efficient routing to a candidate leaf centroid and fast CF updates as the trajectory grows.

An important point follows directly from the criterion: mdBIRCH controls centroid-based average spread (i.e., average RMSD-to-centroid), rather than enforcing a hard per-frame radius cutoff. As a result, individual frames in a cluster can sometimes lie farther than *τ* from the centroid even though the cluster, as a whole, satisfies the specified bound. We return to this point when discussing within-cluster RMSD distributions and RMSD threshold selection.

### 2.5. Streaming updates and order sensitivity

As an online method, mdBIRCH can also be inherently sensitive to the order of data insertion. Earlier assignments influence the centroids and cluster summaries that later frames encounter. In many MD use cases this can be acceptable (and even desirable) because frames arrive in time order and the goal is to update clustering as simulations extend.

This allows the cluster definitions to evolve with the simulation, in contrast to static batch methods that require a complete trajectory upfront to analyze frames from a global perspective. To the best of our knowledge, every single clustering method used to process MD simulations requires to have access to all of the data to be processed, before starting the clustering. mdBIRCH is unique in that it does not need to “see” all the conformations to start clustering them. In cases where order-invariance is important in offline analysis, a straightforward mitigation is to perform multiple runs with different ordering before streaming insertion. In the Results, we quantify order sensitivity under randomized permutations to assess the stability of mdBIRCH clustering in practice.

## 3. SYSTEMS

### -Heptapeptide

Topology and trajectory files were retrieved from a public GitHub repository.^22,23^ Frames 1-1000 were discarded as equilibration, and the remainder was aligned to its first frame. Clustering included a backbone-only representation (N, Cα, C, O, H) restricted to residues Lys2 to Asp11, excluding terminal residues and all side chains, following Daura, et al.^24^ In total, 6,001 structures were included in the analysis.

### HP35 (Nle/Nle mutant)

The long-timescale trajectory from D.E. Shaw Research was used (305 µs at 360 K; frames saved every 200 ps; ∼1.52 × 10^6^ frames).^25^ Frames prior to the 5000^th^ frame were discarded as equilibration, and the remainder was aligned to the 5000^th^ frame. Clustering used a backbone heavy-atom definition consistent with previous HP35 studies: N of residue 1; N, Cα, and C of residues 2-34; and N of residue 35. In total, 1,511,042 structures were used for clustering.

Because mdBIRCH enforces a bound on the post-merge centroid-based spread that is directly related to RMSD (Theory Section 2.3), the threshold *τ* can be interpreted in structural terms. To make this interpretation concrete, we use a small set of deliberately perturbed “reference” structures to define physically meaningful RMSD scales. Specifically, we apply controlled rigid-body rotations to selected residue blocks and compute the RMSD of each edited structure relative to the original reference. These RMSD values serve as anchor points for *τ*: running mdBIRCH with *τ* set equal to one of the RMSDs allows clusters whose centroid-based spread is comparable to the structural deviation introduced by that perturbation. In this way, the reference edits provide an intuitive mapping between numerical *τ* values and recognizable structural changes. The exact perturbations (hinges, moved residue ranges, rotation axes/angles, and resulting RMSDs) are listed in Tables S1-S2.

## 4. RESULTS AND DISCUSSION

### 4.1. Effect of the CF-tree capacity on fragmentation

Using BF (defined in Theory Section 2.4), we tested how the CF-tree capacity affects fragmentation and the yield of well-populated clusters before analyzing *τ*-driven resolution.

We performed a parameter sweep on the HP35 system, testing BFs of 50, 100, 500, and 1000 across a range of RMSD thresholds (0 to 10). As shown in Fig. 2, the clustering behavior follows the same general trends across all BFs, suggesting that the RMSD threshold remains the primary determinant of clustering resolution. However, we observed a clear improvement in data consolidation when using a higher BF.

**Figure 2.**
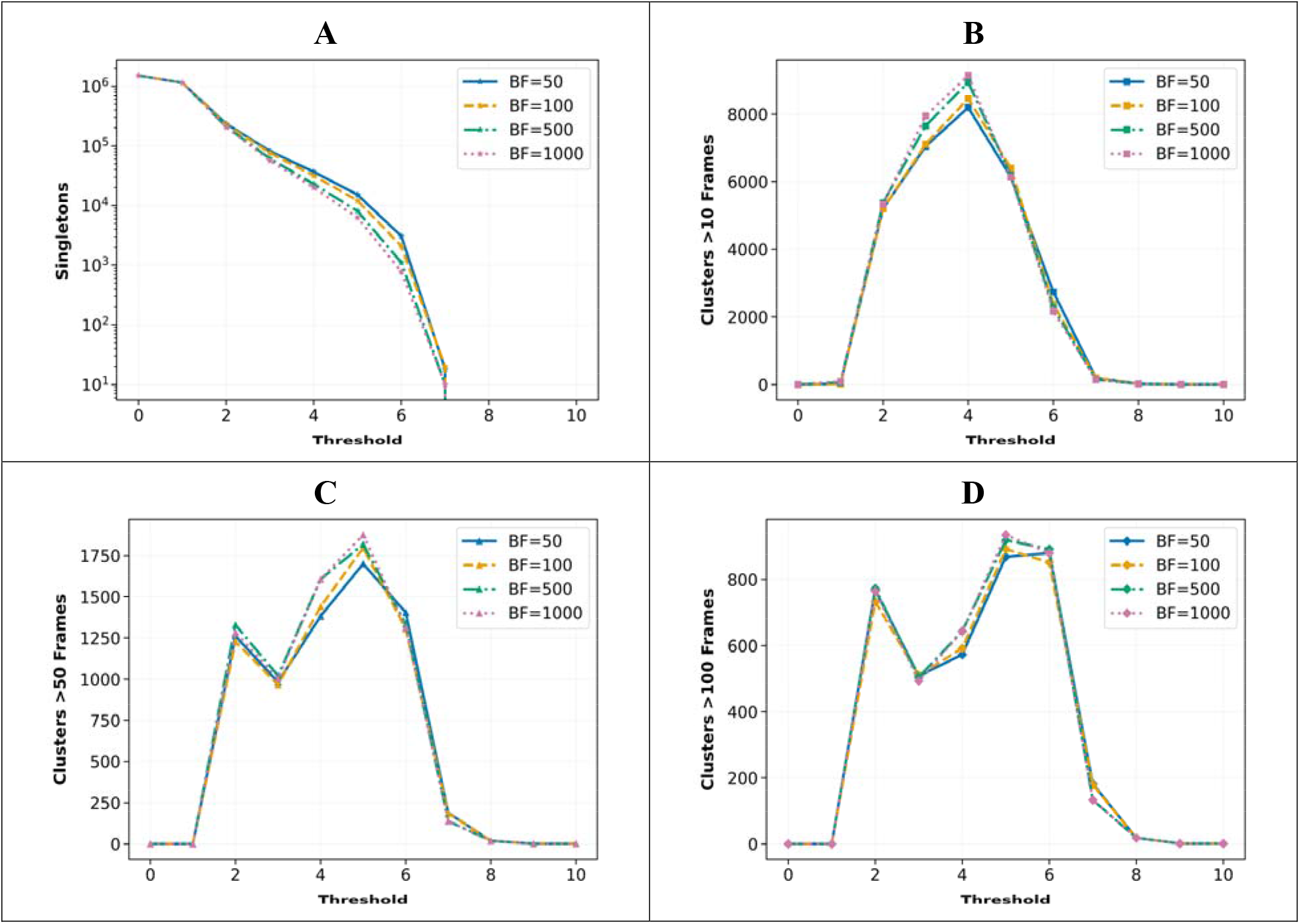
Influence of Branching Factor (BF) on clustering results for the HP35 system. (A) The number of singleton clusters (log scale) versus threshold. (B-D) The number of clusters with >10 frames (B), >50 frames (C), and >100 frames (D) versus threshold.

As presented in Fig. 2A, increasing the BF consistently lowers the number of singleton clusters (clusters containing only one frame) across all thresholds. The high prevalence of singletons at lower BFs suggests that a smaller node capacity can cause more frequent splits and a deeper CF-tree with more, smaller leaf microclusters. Under greedy nearest-centroid insertion, this finer partitioning increases the likelihood that incoming frames form new microclusters rather than merging into existing nearby ones, thereby increasing fragmentation.

Crucially, this reduction in fragmentation translates directly into a higher yield of highly populated clusters. Fig. 2B, 2C, and 2D demonstrate that the run with BF=1000 consistently identifies the highest number of clusters with meaningful populations (frames >10, >50, >100, respectively). This confirms that the higher BF is effectively assigning what would have been singletons into more populated clusters.

Moreover, while a larger BF theoretically increases the computational cost of node updates (as more comparisons are required per level), we found this overhead to be negligible in practice compared to the improvements in clustering quality. Consequently, we fixed BF=1000 for all subsequent analyses in an attempt to limit the cluster behavior only by the physically interpretable threshold, rather than by the capacity of the CF-tree.

### 4.2. Selecting threshold in practice

mdBIRCH exposes a single tolerance in RMSD that governs structural granularity. Because the relevant scales differ across systems and atom selections, we use two complementary strategies: (i) RMSD-anchored operating points obtained from controlled modifications to a representative frame and (ii) blind sweep of the thresholds to visualize how coarseness evolves across a broader range of *τ*.

#### 4.2.1. RMSD-anchored threshold from structural modifications

Using the edited reference structures in Fig. 3, we treat the resulting RMSDs as physically interpretable operating points. We then ran mdBIRCH at each anchored to assess how clustering resolution changes across meaningful structural perturbations.

**Figure 3.**
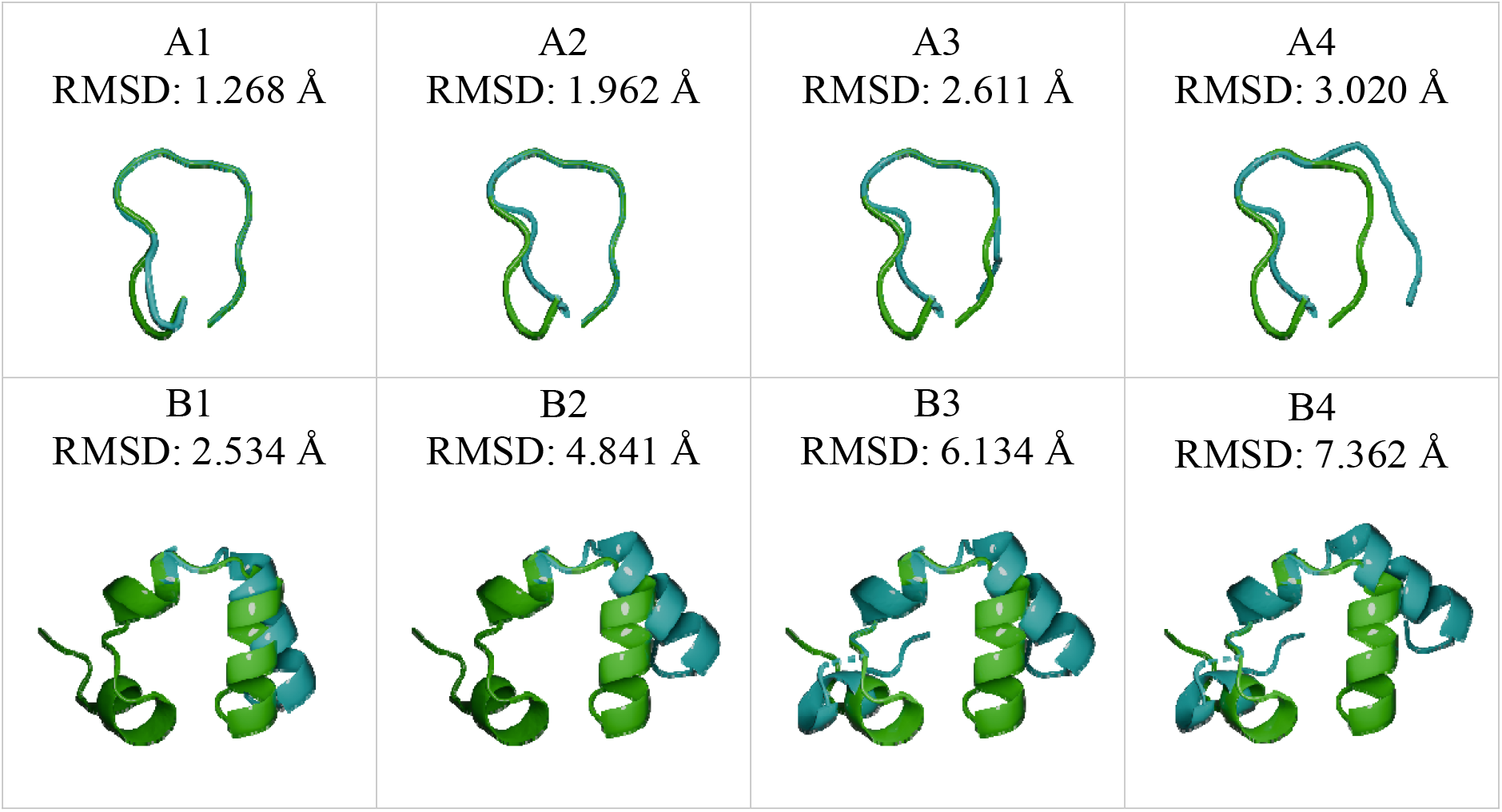
Edited reference overlays used to define RMSD-anchored thresholds. (A) - heptapeptide edits (A1–A4) and (B) HP35 edits (B1–B4). The reference structure is shown in green and edited structures in cyan; titles report the RMSD to the reference (Å) computed using the same atom selection used for clustering.

As *τ* increases, both systems show expected consolidation of states, but on different system-specific scales. For the *β*-heptapeptide (Table 1), raising this *τ* from 1.268 to 3.020 Å collapses the total number of clusters from 658 to 5. In parallel, the number of well-occupied clusters converges to the same five dominant groups: the counts of clusters with >10/>50/>100 frames all become 5 when *τ* is 3.020 Å. The fraction of frames contained in the ten largest clusters also increases monotonically, indicating that at the highest threshold, essentially all the frames are captured by a handful of states.

**Table 1.** Clustering summary at RMSD-anchored thresholds for the -heptapeptide system.

| Threshold<br>( Å ) | Number of<br>Clusters | Clusters<br>>10 Frames | Clusters<br>>50 Frames | Clusters<br>>100 Frames | % Top 10<br>Clusters |
| --- | --- | --- | --- | --- | --- |
| 1.268 | 658 | 124 | 11 | 6 | 26.65 |
| 1.962 | 156 | 110 | 36 | 12 | 35.88 |
| 2.611 | 19 | 18 | 17 | 16 | 80.47 |
| 3.020 | 5 | 5 | 5 | 5 | 100.00 |

For HP35 (Table 2) the absolute cluster counts are larger and the effective range is also shifted upwards. Increasing from 2.534 to 7.362 Å reduces the total number of clusters from 181,344 to 53, while the top ten coverage increases from 20.42% to 81.83%. Notably, the number of moderately populated clusters is highest at intermediate thresholds before collapsing at the largest *τ*, reflecting an intermediate regime where mdBIRCH consolidates many small microclusters into fewer groups without yet merging the ensemble into a handful of very broad states. Together, these results illustrate the expected trade-off: small *τ* values resolve fine-grained conformational structures, whereas larger *τ* yield fewer, higher-occupancy states.

**Table 2.**
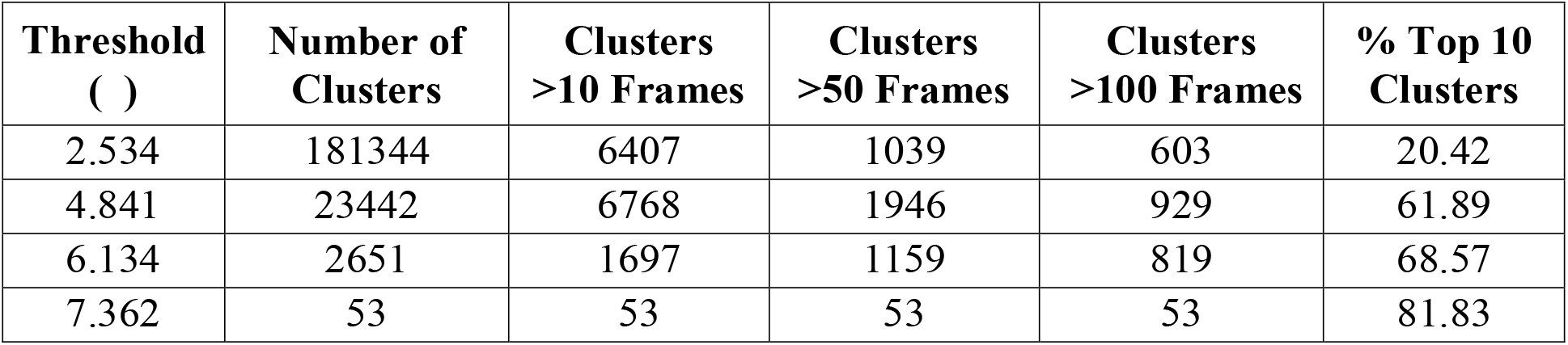
Clustering summary at RMSD-anchored thresholds for the HP35 system.

We next examined the RMSD-to-centroid distributions for the largest clusters at each anchored threshold. Figs. 4 and 5 show the kernel density estimates of RMSD to the cluster centroid for the five largest clusters. For the *β*-heptapeptide (Fig. 4), ncreasing *τ* shifts the distribution to higher RMSDs and generally broadens the tails, consistent with a looser acceptance bound. At *τ* =1.268 Å, dominant modes are concentrated below ∼1 Å with limited tails, reflecting fine-grained clusters. At intermediate thresholds, modes shift forward ∼1-2.5 Å and tails lengthen as nearby conformations consolidate. At *τ* =3.020 Å, only a few high occupancy clusters remain; their densities are broader and partially overlapping, as expected for coarser groupings.

**Figure 4.**
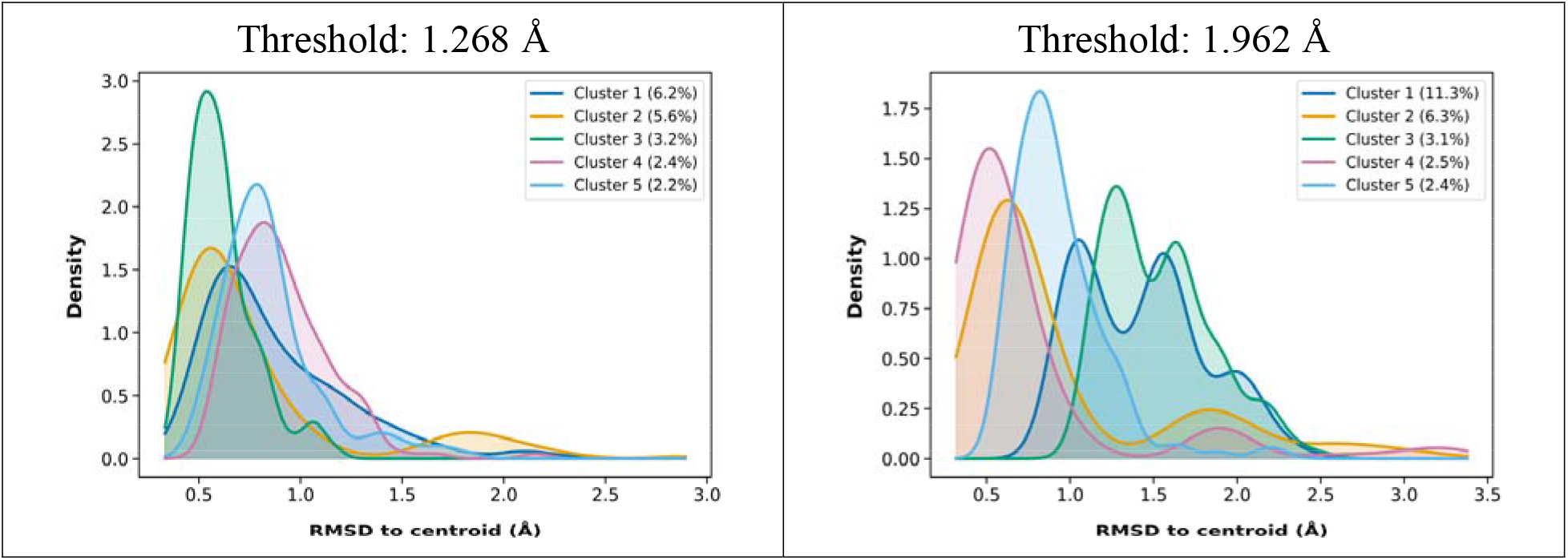
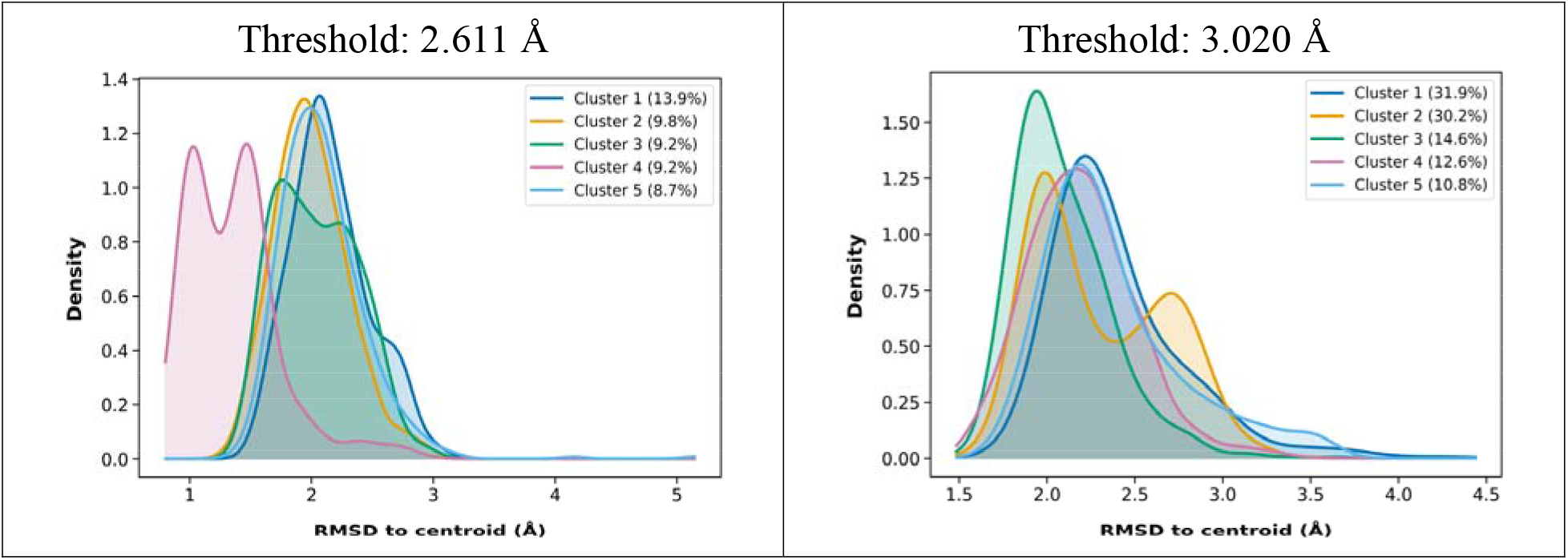
Per-cluster RMSD-to-centroid distributions at RMSD-anchored thresholds for the *β*-heptapeptide system. Curves show kernel density estimates of RMSD to the cluster centroid for the five largest clusters; legends report cluster populations as percentages of the trajectory frames.

**Figure 5.**
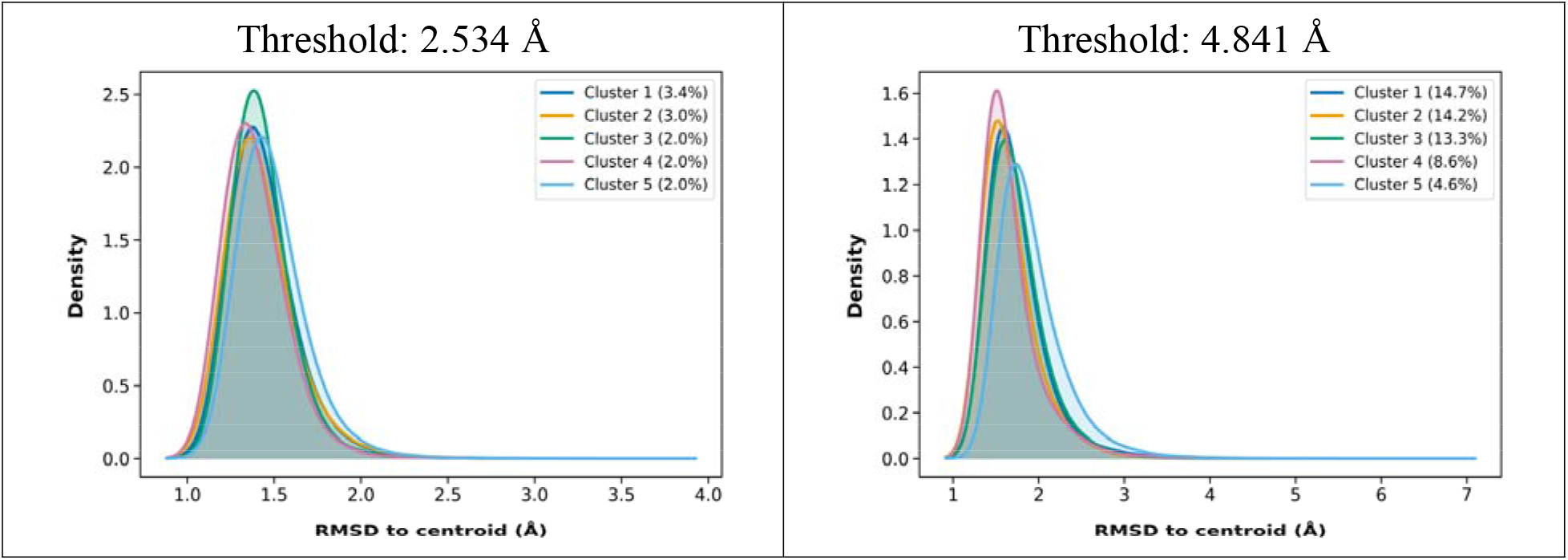
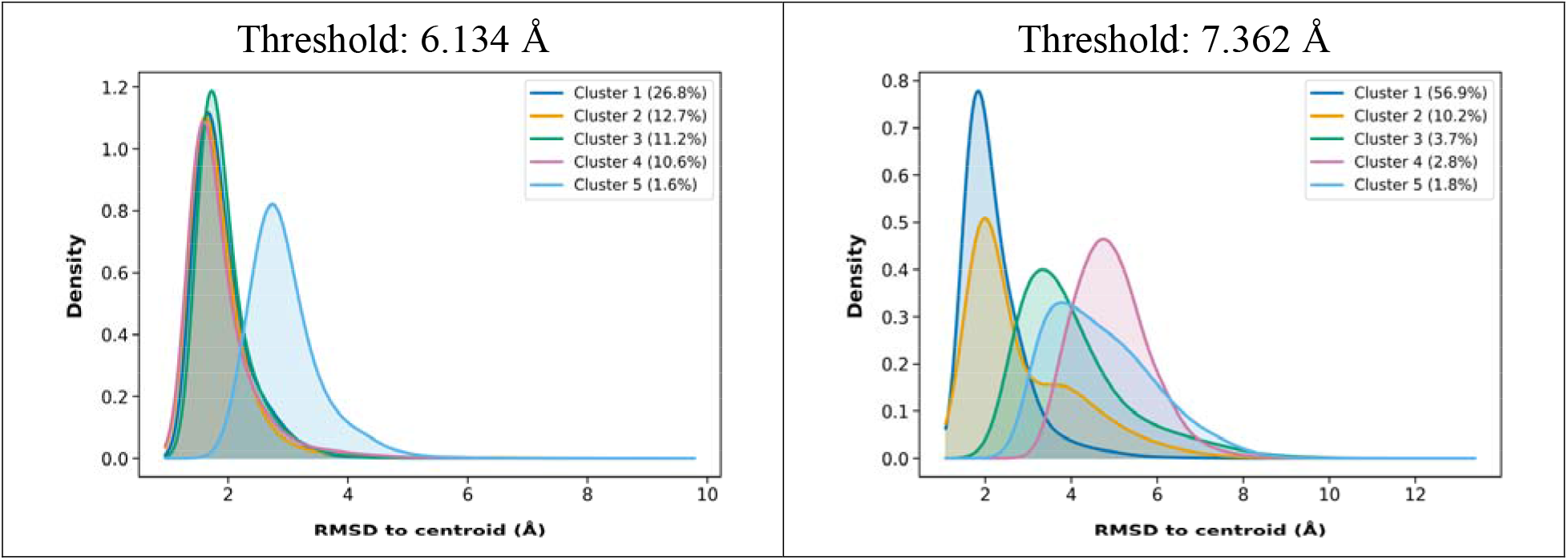
Per-cluster RMSD-to-centroid distributions at RMSD-anchored thresholds for the HP35 system. Curves show kernel density estimates of RMSD to the cluster centroid for the five largest clusters; legends report the cluster populations as percentages of the trajectory frames.

HP35 (Fig. 5) shows the same qualitative behavior on a higher RMSD scale: at the smaller *τ*, clusters exhibit narrow peaks near ∼1.5 Å; at intermediate *τ*, peaks shift upward and tails become heavier as clusters absorb more conformational variability; and at the largest *τ*, a small number of dominant clusters remain with broad densities.

We emphasize here that individual RMSD-to-centroid values can exceed *τ*, because mdBIRCH controls a centroid-based average post-merge spread rather than enforcing a strict maximum-distance cutoff. The observed smooth growth of the modal RMSD with the *τ* is consistent with that guarantee.

#### 4.2.2. Blind threshold sweeps

To complement the RMSD-anchored operating points, we performed blind threshold sweeps over fixed RMSD increments for each system. This protocol maps how clustering granularity evolves across a broad *τ* range, without assuming a reference edit.

Across both systems, the blind sweeps exhibit the same qualitative behavior observed for the anchored thresholds: increasing *τ* monotonically reduces the total number of clusters and increases the fraction of frames captured by the largest clusters, reflecting progressive consolidation of states. However, the *τ* scale at which consolidation becomes pronounced is strongly system dependent. For the *β*-Heptapeptide (Table 3), consolidation occurs at comparatively low thresholds: the ensemble transitions from thousands of small clusters at *τ* ∼ 1 Å to tens of clusters by 2.5 Å and to a small number of dominant clusters by 3 Å. In contrast, HP35 (Table 4) requires substantially higher values to reach comparable consolidation, reflecting broader conformational variability under the alignment and atom selection.

**Table 3.** Clustering summary at thresholds from blind *τ* sweep for the *β*-heptapeptide system.

| Threshold<br>($\tau$) | Total<br>Clusters | Clusters<br>>10 Frames | Clusters<br>>50 Frames | Clusters<br>>100 Frames | % Top 10<br>Clusters |
| --- | --- | --- | --- | --- | --- |
| 0 | 6001 | 0 | 0 | 0 | 0.17 |
| 0.5 | 3336 | 43 | 3 | 0 | 7.08 |
| 1 | 1115 | 81 | 9 | 6 | 22.66 |
| 1.5 | 426 | 159 | 12 | 7 | 27.45 |
| 2 | 137 | 99 | 37 | 14 | 36.69 |
| 2.5 | 26 | 25 | 23 | 20 | 68.44 |
| 3 | 5 | 5 | 5 | 5 | 100.00 |
| 3.5 | 1 | 1 | 1 | 1 | 100.00 |
| 4 | 1 | 1 | 1 | 1 | 100.00 |
| 4.5 | 1 | 1 | 1 | 1 | 100.00 |
| 5 | 1 | 1 | 1 | 1 | 100.00 |

**Table 4.** Clustering summary at thresholds from blind *τ* sweep for the HP35 system.

| **Threshold**<br>**$\tau$** | **Total**<br>**Clusters** | **Clusters**<br>**>10 Frames** | **Clusters**<br>**>50 Frames** | **Clusters**<br>**>100 Frames** | **% Top 10**<br>**Clusters** |
| --- | --- | --- | --- | --- | --- |
| 0 | 1511042 | 0 | 0 | 0 | 0.00 |
| 1 | 1287956 | 88 | 0 | 0 | 0.01 |
| 2 | 305608 | 5331 | 1283 | 765 | 8.40 |
| 3 | 123288 | 7950 | 1004 | 493 | 36.25 |
| 4 | 53984 | 9141 | 1606 | 643 | 51.08 |
| 5 | 19548 | 6136 | 1875 | 935 | 62.37 |
| 6 | 3787 | 2170 | 1314 | 880 | 67.01 |
| 7 | 153 | 141 | 138 | 133 | 73.75 |
| 8 | 18 | 18 | 18 | 18 | 91.41 |
| 9 | 1 | 1 | 1 | 1 | 100.00 |
| 10 | 1 | 1 | 1 | 1 | 100.00 |

The occupancy filters help distinguish regimes that are often most useful in practice. Very small *τ* values produce extensive fragmentation, with many clusters failing to reach even modest population cutoffs. At intermediate *τ*, the number of well-occupied clusters increases while the total count decreases, indicating that mdBIRCH is consolidating small groups into a set of more interpretable populated clusters. At sufficiently large *τ*, both systems collapse into a few dominant clusters (Tables 3-4), signaling an over-merged regime that is typically too coarse for downstream structural analysis.

Taken together, the anchored runs and blind sweeps provide a practical, reproducible way to select *τ* . From an anchored set of candidate values, a simple heuristic is to choose *τ* near the point where the clustering yields well-occupied clusters and where coverage increases sharply, since these anchors define system-specific, physically interpretable operating points for *τ*. Blind sweeps then serve as a complementary check to (i) verify that the chosen *τ* lies in a sensible regime (neither excessively fragmented nor over-merged) and (ii) identify a reasonable operating range when no prior anchor is available. In practice, we therefore use RMSD-anchored runs as the primary way to set *τ* and treat blind sweeps as an optional diagnostic to confirm and fine-tune that choice, depending on the analysis goal (e.g., separating major basins versus emphasizing finer substructures).

### 4.3. Effect of data ordering

Building on the order-sensitivity described earlier, we quantify how insertion order affects practical outputs by repeating the HP35 sweep under the original trajectory and three randomized permutations, while keeping all parameters fixed.

Fig. 6 shows that the global resolution trends are preserved under reordering. Across all runs, increasing the *τ* produces the same qualitative behavior as seen before. Thus, while mdBIRCH is not strictly order-invariant, the clustering summaries we use here to choose *τ* are only weakly sensitive to permutations for this system.

**Figure 6.**
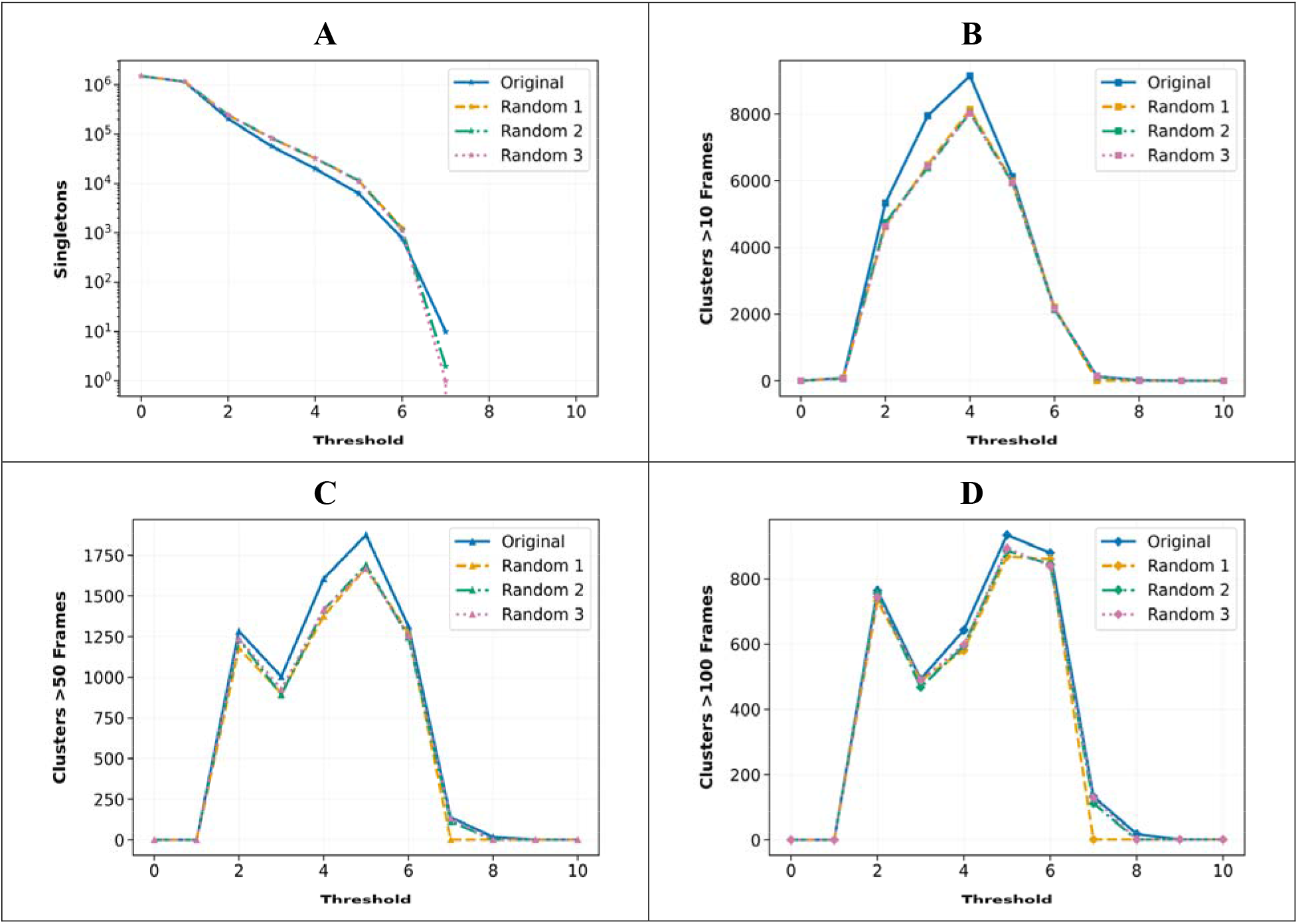
Influence of data ordering on mdBIRCH clustering results for the HP35 system. The original trajectory order (“Original”) is compared to three randomized frame permutations (“Random 1-3”) across RMSD thresholds. (A) The number of singleton clusters (log scale) versus threshold. (B-D) The number of clusters with >10 frames (B), >50 frames (C), and >100 frames (D) versus threshold.

The largest deviation between orderings occurs in the intermediate regimes where the CF-tree is actively balancing “merge” versus “start-new-microcluster” decisions and where small differences in early centroids can propagate to later assignments. Practically, this implies that (a) at very small *τ* (where most points remain isolated) and very large *τ* (where most points consolidate) clustering results are relatively insensitive to ordering, whereas (b) intermediate can show small variability in counts of marginal clusters. This behavior is expected for a streaming clustering method as local greedy decisions are constantly made without access to the full dataset.

Importantly, we highlight here that order dependence is not necessarily undesirable in our intended use case. In an online setting, the natural arrival order is the simulation time order, and mdBIRCH provides a consistent evolving partition that can be updated as frames are produced. For offline analyses where order-invariance is preferred, we recommend two possible mitigations: (i) run multiple permutations, and (ii) shuffle within temporal blocks to preserve locality while reducing strict dependence on arrival order.

### 4.4. Comparison with batch methods

In this section, we compare the dominant clusters identified by mdBIRCH with those obtained from batch clustering workflows to assess their structural correspondence.

Specifically, we compare mdBIRCH’s dominant states against two representative batch pipelines: *k*-means NANI (hereafter referred to as NANI) and HELM. mdBIRCH determines the number of clusters through its physically interpretable tolerance *τ*, while NANI and HELM are typically used in fixed-mode where all frames are allocated into exactly *k* clusters. Because their operations differ, we do not expect strict one-to-one cluster correspondence. Enforcing a *k* can merge conformational basins that mdBIRCH would keep distinct given the threshold, or conversely, split a basin that mdBIRCH would retain as a single state. We therefore compare representative structures of dominant states rather than attempting label-level agreement.

We ran mdBIRCH at *τ*=3 Å and retained only the ten most populated clusters for comparison. For NANI, we used strat_reduced initialization type and generated *k*=10 clusters. For HELM, we started with 60 NANI microclusters and built the hierarchy using the intra linkage and then reported *k*=10 clusters in two variants: with trimming (retaining clusters with MSD<20) and without trimming.

Within each cluster, we selected a medoid, which we defined as the frame with the highest complementary MSD (i.e., the frame whose exclusion produces the largest increase in the set’s MSD). We computed the pairwise RMSD matrix between mdBIRCH medoids and batch method medoids. Heatmaps report these RMSDs where smaller values indicate closer representative structures. These maps are not expected to be diagonal as cluster indices are ordered by population rather than structural correspondence. Agreement is instead reflected by low-RMSD entries anywhere along a row, indicating that a given mdBIRCH dominant state has a close counterpart in the batch partition.

The comparisons indicate that most mdBIRCH clusters have a counterpart in the batch clustering methods, but the apparent correspondence depends on how strongly the batch pipeline enforces compactness. In the HELM (trimmed) variant (Fig. 7), many mdBIRCH medoids find a nearest HELM-trimmed medoid at low RMSD, consistent with both approaches emphasizing structurally tight, physically coherent states. In contrast, HELM without trimming yields substantially larger RMSDs across much of the matrix (Fig. S1), consistent with retaining broader/high-MSD clusters whose representatives can sit far from the compact dominant states emphasized in mdBIRCH at the given threshold. The NANI comparison shows intermediate behavior between the two HELM cases (Fig. S2): several mdBIRCH dominant states map closely to the NANI medoids, while other states exhibit larger RMSDs. This suggests that the fixed *k* constraint here in NANI redistributes structurally heterogeneous regions across a small number of clusters. Repeating the comparison at a slightly coarser tolerance (*τ* =4 Å) produces the same qualitative trends (Figs. S3-S5).

**Figure 7.**
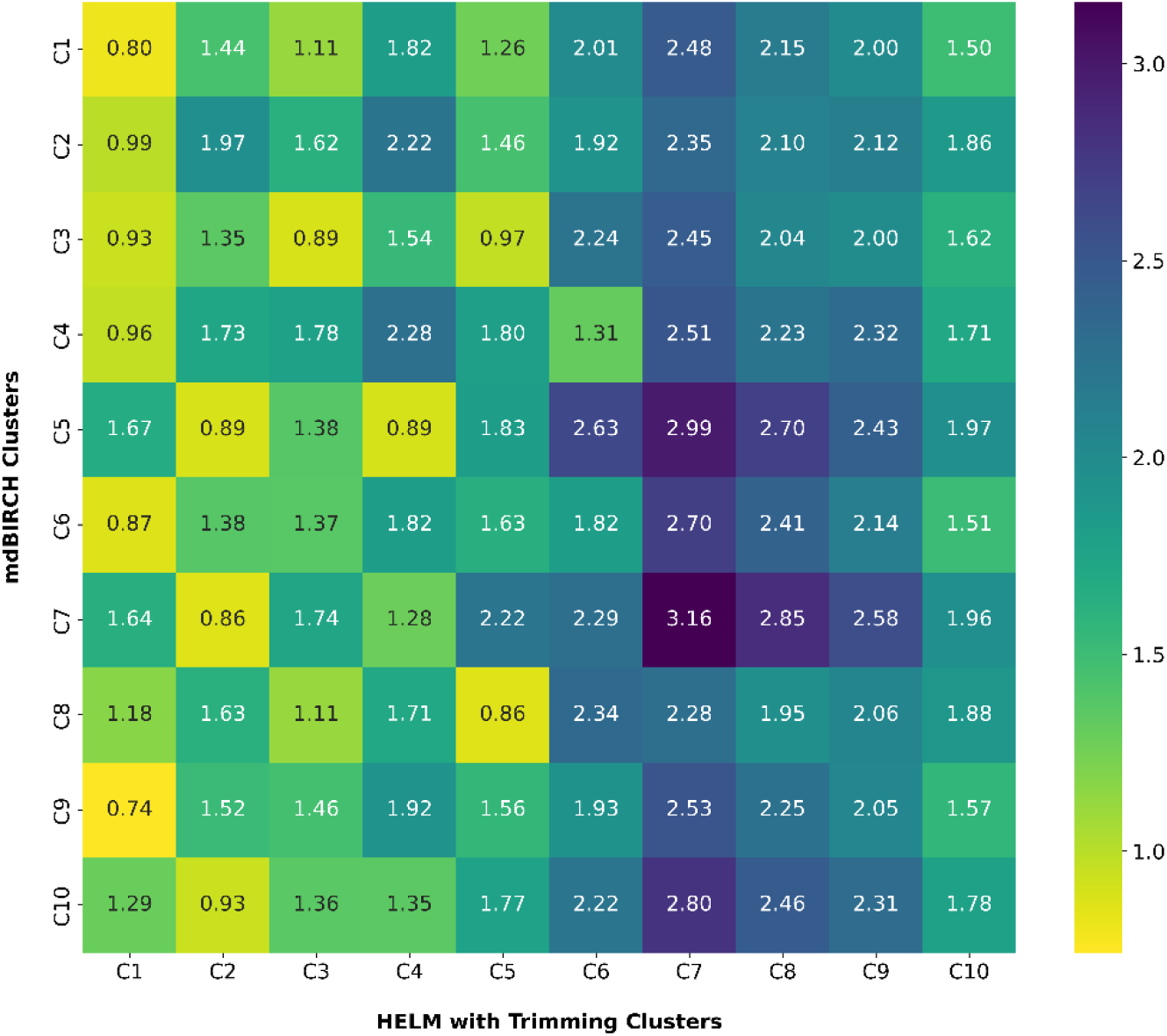
Medoid-to-medoid RMSD matrix comparison on the HP35 system. The heatmap reports pairwise RMSDs (Å) between cluster medoids from mdBIRCH (rows; the 10 most populated clusters, BF=1000, =3 Å) and HELM with trimming (columns; =10, MSD <20, intra linkage). Lighter colors (yellow) indicate lower RMSD and better structural agreement.

However, we highlight here that at the thresholds we used, the full partition of mdBIRCH contains many clusters (>10); an exhaustive cross-method matching would therefore require comparing a much larger set of representatives. To keep the analysis interpretable, we restrict attention to the ten most populated mdBIRCH clusters, which capture the dominant states typically used for visualization and downstream modeling. Consequently, some batch-method clusters may correspond more closely to the less-populated mdBIRCH clusters not included in this top ten subset.

### 4.5. Runtime scaling and practical throughput

To assess mdBIRCH’s computational efficiency, we measured the wall-time as a function of the number of frames in a single pass (BF=1000, *τ*=3). As shown in Fig. 8, the runtime grows smoothly and close to linear up to 300k frames, with some variance across repeated runs. This behavior is consistent with the mdBIRCH implementation; each insertion is routed to the nearest centroid through a CF-tree and performs a single post-merge check, so the number of centroid comparisons is bound by tree capacity and no pairwise distance matrix calculation is made. In practical terms, mdBIRCH processes hundreds of thousands of frames on a single CPU core in seconds. All runtime measurements were performed using a standard Python implementation of mdBIRCH executed on a single CPU core (AMD Ryzen 7 8845HS at 3.80 GHz) with 16 GB RAM. No parallelization or GPU acceleration was used.

**Figure 8.**
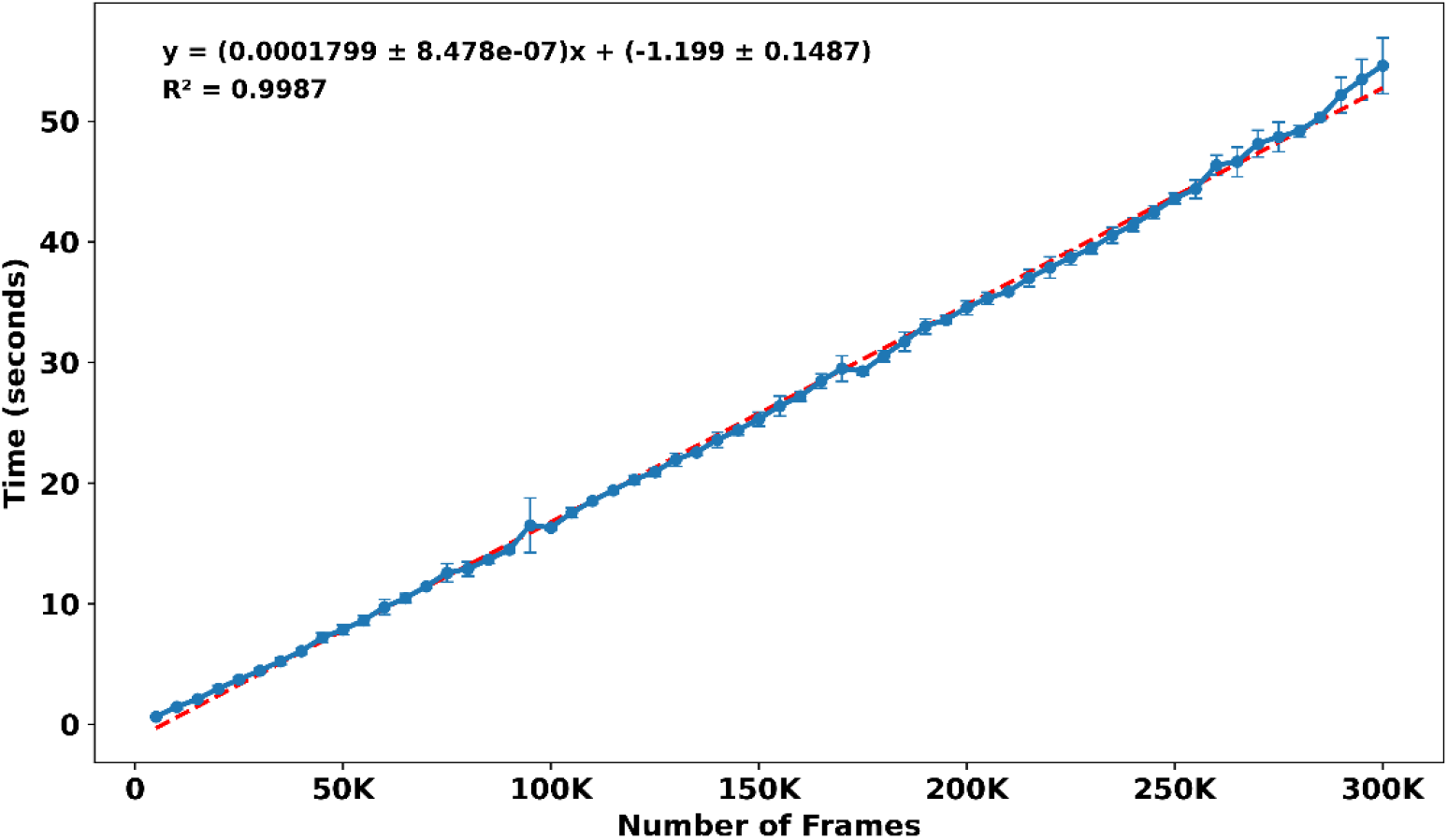
Runtime scaling of mdBIRCH with trajectory length of HP35. Wall-clock time (seconds) versus the number of frames with the HP35 system (5,000 to 300,000 frames; =3 Å; BF=1000; CPU timing). Points show the mean across five repeats; error bars indicate the standard deviation. The dashed red line is a least-squares fit (equation and R^2^ reported on the figure).

We want to highlight that even this impressive performance is the worst-case scenario for mdBIRCH. That is, in this benchmark we illustrated how mdBIRCH would scale if the dataset is processed after the whole MD simulation has been run. In a more practical scenario, mdBIRCH will be coupled to the MD engine, in a way that every new frame generated will pass through the tree before the next MD frame has been saved. This makes mdBIRCH effectively instantaneous, able to provide a full analysis of the trajectory immediately after the last conformation has been produced.

## 5. CONCLUSIONS

We set out to keep online clustering of trajectories simple, scalable, and physically interpretable. mdBIRCH achieves this by combining the BIRCH CF-tree with a merge decision calibrated in RMSD units. Each arriving frame is routed to a candidate leaf-microcluster, and we evaluate the candidate cluster after the hypothetical inclusion of the new frame. The insertion is accepted if the post-merge cluster satisfies the RMSD-calibrated bound implied by the user-chosen threshold ; otherwise, a new microcluster is created. This design provides a single-pass guarantee on centroid-based spread while remaining practical for large-scale analysis.

On both the -heptapeptide and HP35 systems, behaves as an intuitive control for granularity. Using RMSD-anchored operating points from controlled structural edits, together with blind threshold sweeps, we observed consistent and system-specific trends: increasing reduces the total number of clusters, consolidates population into high-occupancy states, and broadens the RMSD-to-centroid distributions within clusters. Importantly, these distributions also clarify the interpretation of the guarantee: mdBIRCH constrains centroid-based average spread rather than the maximum pairwise RMSD, so individual frames may lie beyond . In addition, we found that CF-tree search parameter BF can influence clustering outcomes: increasing the BF reduces fragmentation (e.g., fewer singleton clusters) and improves assignment into well-populated states.

From a computational perspective, mdBIRCH is fast and memory-bounded by construction. Because each insertion requires only tree routing plus a single post-merge check computed from CF summaries, runtime grows smoothly and close to linearly with trajectory on standard CPU hardware, enabling dense threshold scans and routine sensitivity checks that are difficult to justify with quadratic pairwise methods. We also show that while mdBIRCH can exhibit minimal order dependence, as expected for streaming algorithms, this behavior aligns with its intended use case when frames arrive in simulation time order. Furthermore, we note here that like other pure geometric clustering approaches, mdBIRCH does not inherently account for kinetic criteria: it identifies geometrically compact structural states, which may align with kinetically metastable states but are not guaranteed to do so (which is a question that will be pursued elsewhere).^27^

In future work, we plan to explore tighter integration of mdBIRCH with MD engines, enabling clustering decisions to be performed in real time as frames are generated. The incremental nature of mdBIRCH makes it well suited for coupling with adaptive sampling workflows and on-the-fly analyses pipelines. Such integration would allow immediate identification of emerging structural states and could facilitate efficient feedback-driven simulation strategies.

Overall, mdBIRCH combines efficient, incremental partitioning with a physically interpretable tolerance, making it well suited for long simulations that are extended over time or for ensembles that are appended as new segments arrive. Remarkably, all of this can be achieved at the lowest time and memory cost, with essentially no overhead while the simulation is running. With no need to discard precious MD frames, and the possibility of having full clustering results immediately after the simulation concluded, mdBIRCH offers key advantages over more traditional methods.

## ACKNOWLEDGEMENTS

We thank support from the National Institute of General Medical Sciences of the National Institutes of Health under award number R35GM150620.

## DATA AND SOFTWARE AVAILABILITY

The mdBIRCH code can be found here: https://github.com/mqcomplab/MDANCE.

### Supporting Information

Derivation of the RMSD-calibrated nucleation and merge criterion; details of the structural perturbations used to define RMSD-anchored thresholds; and additional comparisons with batch clustering methods including medoid-to-medoid RMSD matrices between mdBIRCH and HELM or *k*-means NANI clustering results.

## TOC Graphic

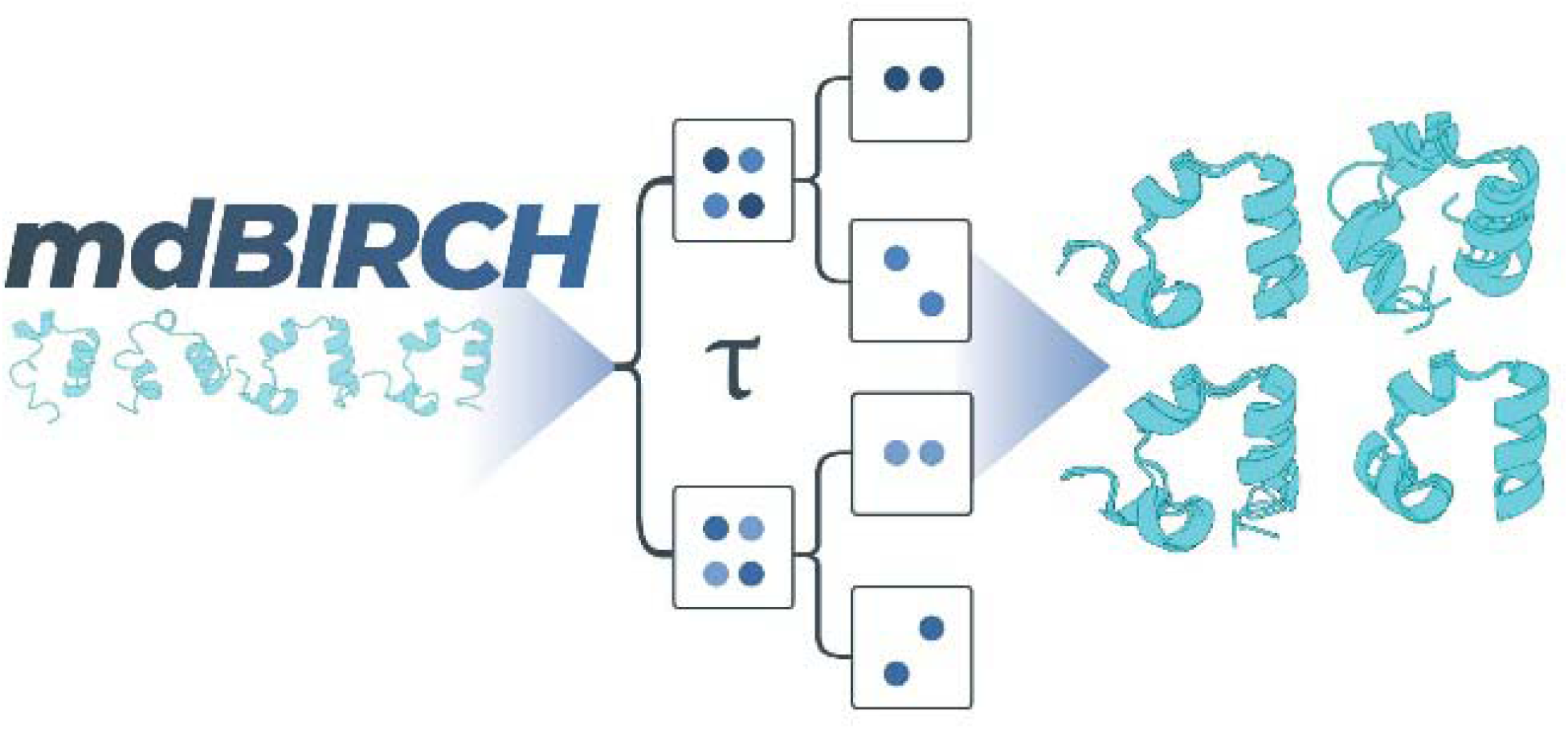

