## Supplementary material for "mdBIRCH for Fast, Scalable, Online Clustering of Molecular Dynamics Trajectories": SI

**SUPPORTING INFORMATION**

**Derivation of the RMSD-calibrated nucleation and merge criterion**

This section describes the relationship between the cluster’s squared radius ($R^{2}$) and the pairwise RMSD for the case of a two-frame cluster. This relationship forms the basis for the nucleation and the merge criterion in mdBIRCH.

A cluster $j$ containing $N_{j}$ frames is summarized by its Cluster Feature ($\text{CF}_{j}$) in Eq. (S1):

$$\begin{aligned} \text{CF}_{j}=\left( N_{j},{LS}_{j},{SS}_{j} \right), \#\left( S1 \right) \end{aligned}$$

where ${LS}_{j}$ is the vector sum of all frames in the cluster and ${SS}_{j}$ is the scalar sum of the squared Euclidean norms of the frames.

The $R_{j}^{2}$ is the mean squared radius about the centroid $c_{j}={LS}_{j}/N_{j}$. As shown in Eq. (6), this is calculated via Eq. (S2):

$$\begin{aligned} R_{j}^{2}=\frac{{SS}_{j}}{N_{j}}-\frac{\left\| {LS}_{j} \right\|^{2}}{N_{j}^{2}}.\#\left( S2 \right) \end{aligned}$$

The pairwise RMSD between two aligned frames, $x_{1}$ and $x_{2}$ (represented as vectors in D dimensions), is defined in Eq. (S3):

$$\begin{aligned} \text{RMSD}\left( x_{1},x_{2} \right)=\sqrt{\frac{1}{D}\sum_{i=1}^{D} \left( x_{1i}-x_{2i} \right)^{2}}=\sqrt{\frac{1}{D}\left\| x_{1}-x_{2} \right\|^{2}}.\#(S3)\# \end{aligned}$$

Squaring both sides gives Eq. (S4):

$$\begin{aligned} \text{RMSD}\left( x_{1},x_{2} \right)^{2}=\frac{1}{D}\left\| x_{1}-x_{2} \right\|^{2}.\#\left( S4 \right) \end{aligned}$$

Rearranging gives the direct relationship between the squared Euclidean distance and squared RMSD shown in Eq. (S5):

$$\begin{aligned} \left\| x_{1}-x_{2} \right\|^{2}=D\cdot\text{RMSD}\left( x_{1},x_{2} \right)^{2}. \# \left( S5 \right) \end{aligned}$$

*Two-Frame Cluster*

Consider the nucleation of a new cluster from two frames, $x_{1}$ and $x_{2}$. For this new cluster, the summaries are given by Eq. (S6):

$$\begin{aligned} N=2, LS=x_{1}+x_{2}, SS=\left\| x_{1} \right\|^{2}+\left\| x_{2} \right\|^{2}.\#\left( S6 \right) \end{aligned}$$

We can now calculate $R^{2}$ using Eq. (S2):

$$R^{2}=\frac{SS}{N}-\frac{\left\| LS \right\|^{2}}{N^{2}}$$

$$R^{2}=\frac{\left\| x_{1} \right\|^{2}+\left\| x_{2} \right\|^{2}}{2}-\frac{\left\| x_{1}+x_{2} \right\|^{2}}{2^{2}}$$

$$R^{2}=\frac{\left\| x_{1} \right\|^{2}+\left\| x_{2} \right\|^{2}}{2}-\frac{\left\| x_{1} \right\|^{2}+\left\| x_{2} \right\|^{2}+2(x_{1}\cdot x_{2})}{4}$$

$$R^{2}=\frac{{2\left\| x_{1} \right\|}^{2}+\left\| x_{2} \right\|^{2}}{4}-\frac{\left\| x_{1} \right\|^{2}+\left\| x_{2} \right\|^{2}+2(x_{1}\cdot x_{2})}{4}$$

$$R^{2}=\frac{1}{4}(\left( {2\left\| x_{1} \right\|}^{2}+\left\| x_{2} \right\|^{2} \right)-(\left\| x_{1} \right\|^{2}+\left\| x_{2} \right\|^{2}+2\left( x_{1}\cdot x_{2} \right)))$$

$$R^{2}=\frac{1}{4}(\left\| x_{1} \right\|^{2}+\left\| x_{2} \right\|^{2}-2\left( x_{1}\cdot x_{2} \right))$$

We recognize the final term as the expansion of a squared difference; hence, we get Eq. (S7):

$$\begin{aligned} R^{2}=\frac{1}{4}\left( \left\| x_{1}-x_{2} \right\|^{2} \right).\#\left( S7 \right) \end{aligned}$$

This intermediate result shows that the cluster’s squared radius ($R^{2}$) for two frames is exactly one quarter of their squared Euclidean distance.

*Conversion to RMSD units*

Finally, we substitute the definition from Eq. (S4) into Eq. (S7) to get Eq. (S8) (which corresponds to Eq. (10) in the main manuscript):

$$\begin{aligned} R^{2}=\frac{1}{4}D\cdot\text{RMSD}\left( x_{1},x_{2} \right)^{2}. \#\left( S8 \right) \end{aligned}$$

**Structural edits used to define RMSD-anchored thresholds**

To support the selection of RMSD-anchored operating points for the threshold, we provide here details of the structural perturbations used in Section 4.2.1. For each system, the reference structure was taken as the final frame of the corresponding MD simulation, and each modification was generated by applying one or two rigid rotations to a specific residue block about a hinge C$\alpha$ atom in PyMOL. For each rotation, the hinge was set as the rotation pivot, followed by a rotation of the moving selection. RMSDs were then computed relative to the unmodified reference under the same atom selection used for clustering. Tables S1-S2 list, for each modification, the hinge atom, moving segment, rotation axis and angle(s), and the resulting RMSD that was used as a candidate threshold value.

**Table S1.** Controlled structural modifications for the $\beta$-heptapeptide system.

| **ID** | **Operation** | **Hinge  (pivot atom)** | **Moving Residues** | **Axis, Angle** | **Resulting RMSD (Å)** |
| --- | --- | --- | --- | --- | --- |
| A1 | single rotation | resi 4, CA | 2-4 | [-2, 1, -1], -75° | 1.268 |
| A2 | single rotation | resi 5, CA | 2-5 | [1, 1, -2], 30° | 1.962 |
| A3 | two rotations | resi 5, CA resi 9, CA | 2-5  9-12 | [1, 1, -2], 30°  [1, 1, -2], 45° | 2.611 |
| A4 | two rotations | resi 5, CA resi 7, CA | 2-5  7-12 | [1, 1, -2], 30°  [1, 3, 2], -30° | 3.020 |

**Table S2.** Controlled structural modifications for the HP35 system.

| **ID** | **Operation** | **Hinge  (pivot atom)** | **Moving Residues** | **Axis, Angle** | **Resulting RMSD (Å)** |
| --- | --- | --- | --- | --- | --- |
| B1 | single rotation | resi 20, CA | resi 21-35 | [0, 1, 1], 30° | 2.534 |
| B2 | single rotation | resi 20, CA | resi 21-35 | [0, 2, 4], 45° | 4.841 |
| B3 | two rotations | resi 20, CA resi 12, CA | resi 21-35  resi 1-11 | [0, 2, 4], 45°  [-2, -4, -4], 90° | 6.134 |
| B4 | two rotations | resi 20, CA resi 12, CA | resi 21-35  resi 1-11 | [0, 2, 4], 60°  [-2, -4, -4], 90° | 7.362 |

**Comparison with batch methods**

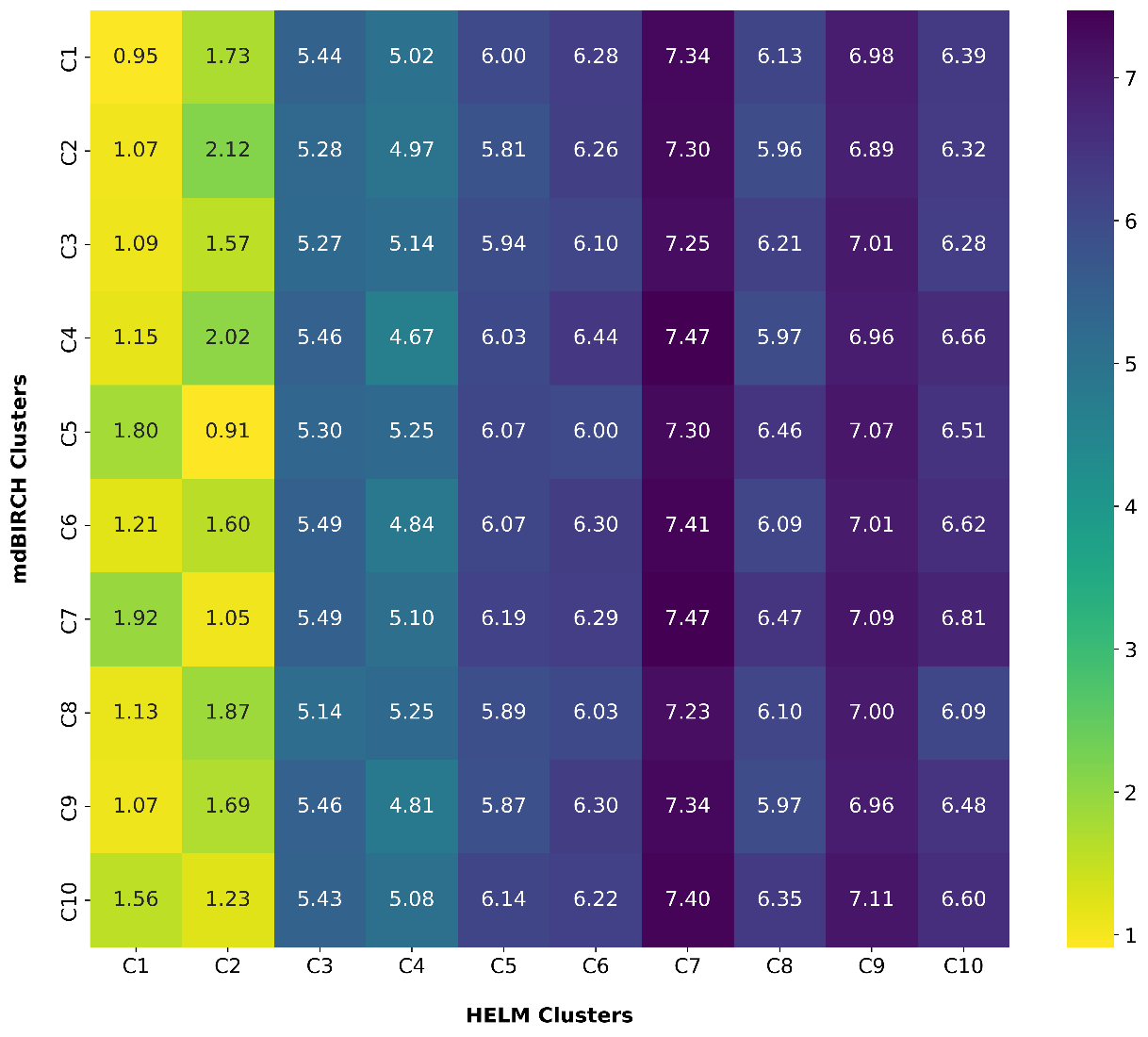

**Figure S1.** Medoid-to-medoid RMSD matrix comparison on the HP35 system. The heatmap reports pairwise RMSDs (Å) between cluster medoids from mdBIRCH (rows; the 10 most populated clusters, BF=1000, $\tau$=3 Å) and HELM with no trimming (columns; $k$=10, intra linkage). Lighter colors (yellow) indicate lower RMSD and better structural agreement.

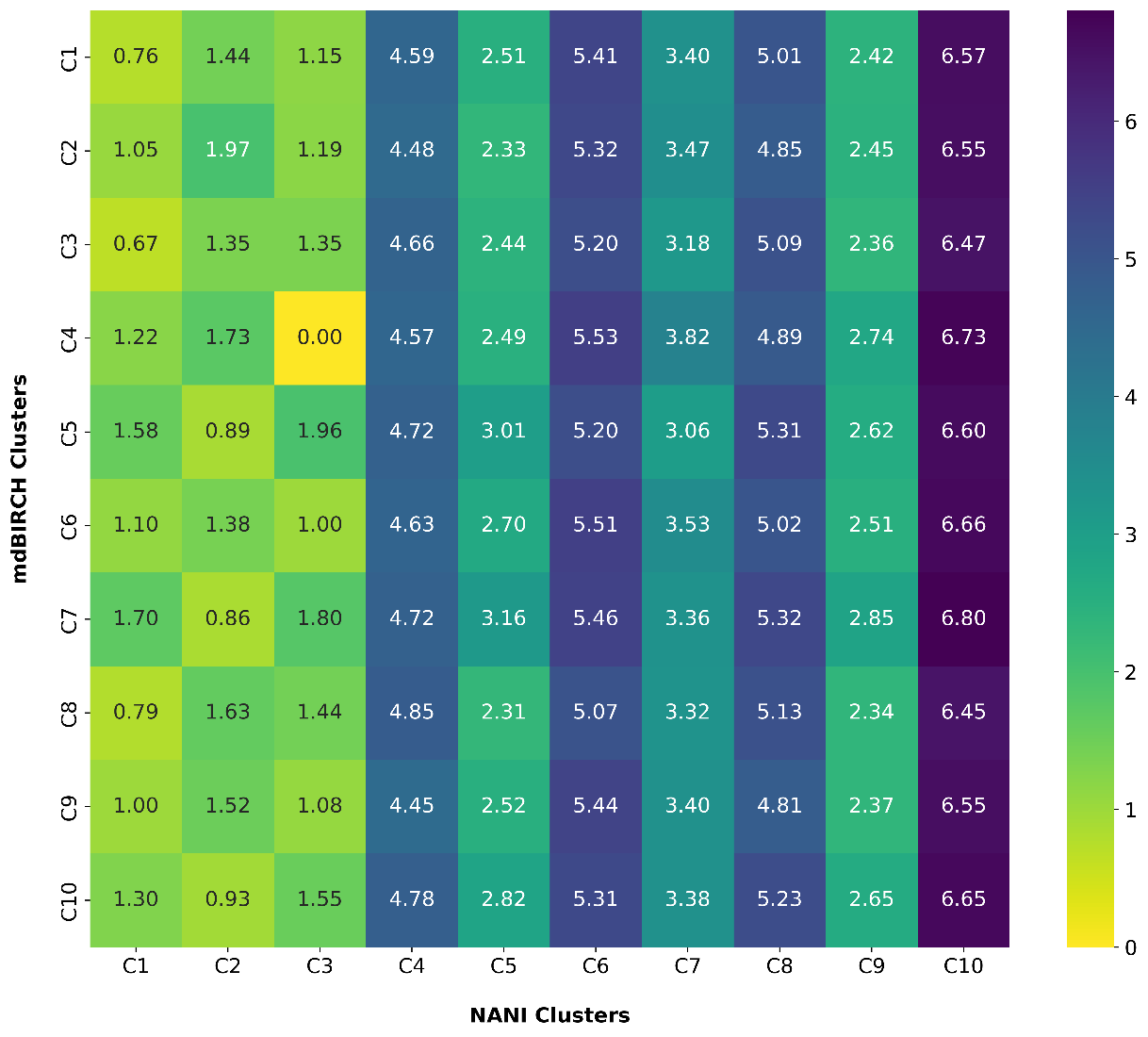

**Figure S2.** Medoid-to-medoid RMSD matrix comparison on the HP35 system. The heatmap reports pairwise RMSDs (Å) between cluster medoids from mdBIRCH (rows; the 10 most populated clusters, BF=1000, $\tau$=3 Å) and *k*-means NANI (columns; $k$=10, strat_reduced initialization type). Lighter colors (yellow) indicate lower RMSD and better structural agreement.

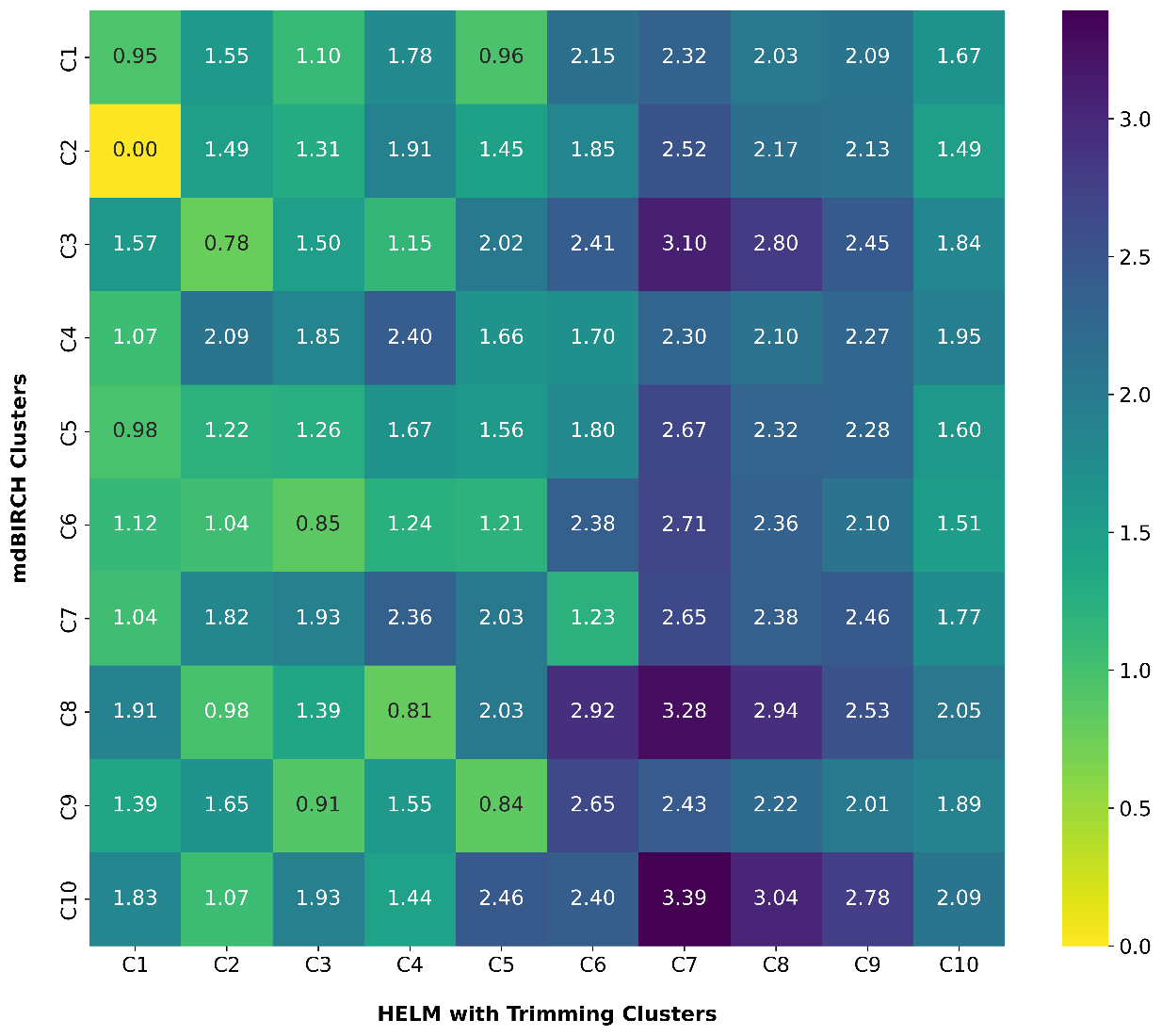

**Figure S3.** Medoid-to-medoid RMSD matrix comparison on the HP35 system. The heatmap reports pairwise RMSDs (Å) between cluster medoids from mdBIRCH (rows; the 10 most populated clusters, BF=1000, $\tau$=4 Å) and HELM with trimming (columns; $k$=10, using clusters with MSD <20, intra linkage). Lighter colors (yellow) indicate lower RMSD and better structural agreement.

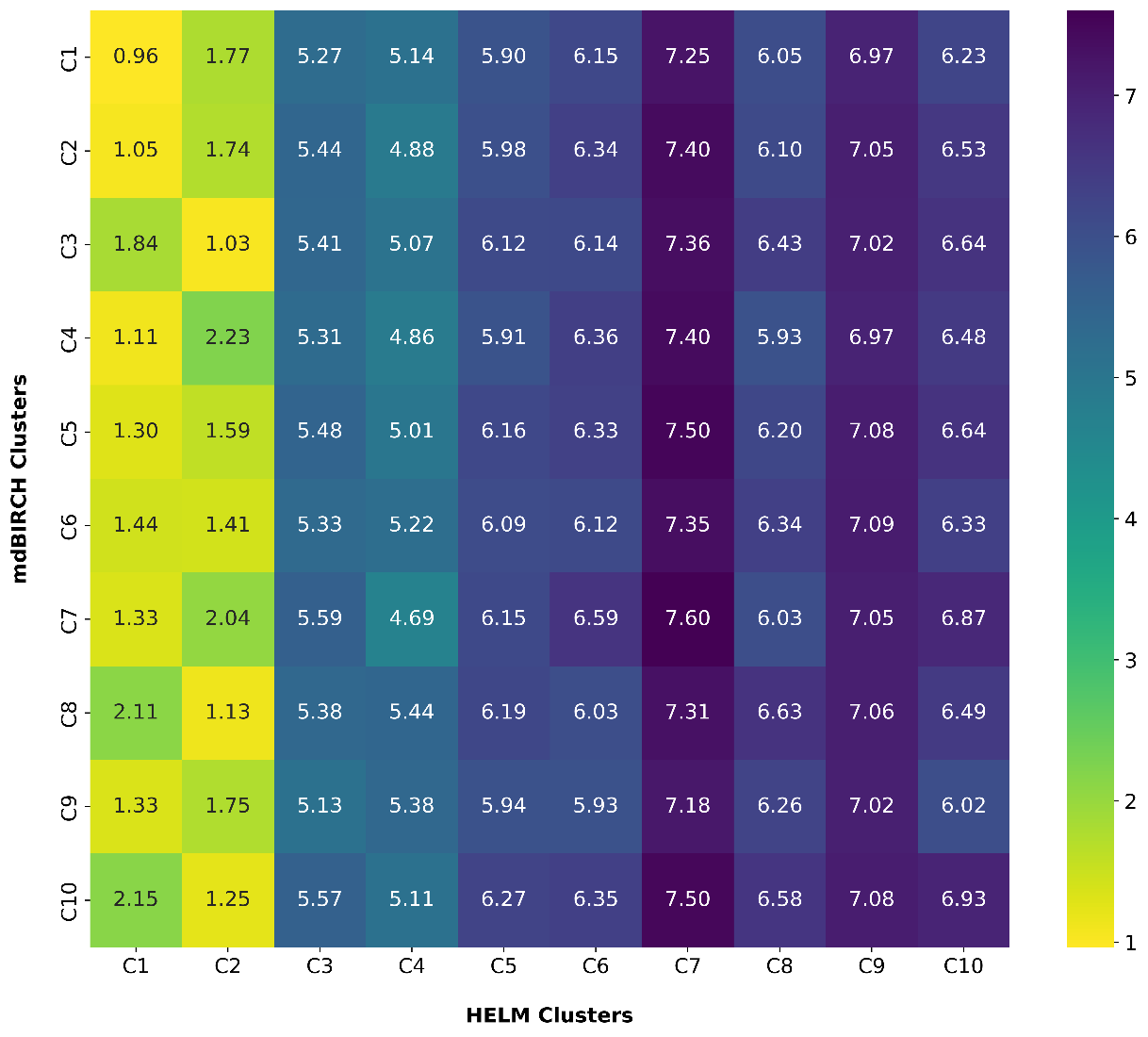

**Figure S4.** Medoid-to-medoid RMSD matrix comparison on the HP35 system. The heatmap reports pairwise RMSDs (Å) between cluster medoids from mdBIRCH (rows; the 10 most populated clusters, BF=1000, $\tau$=4 Å) and HELM with no trimming (columns; $k$=10, intra linkage). Lighter colors (yellow) indicate lower RMSD and better structural agreement.

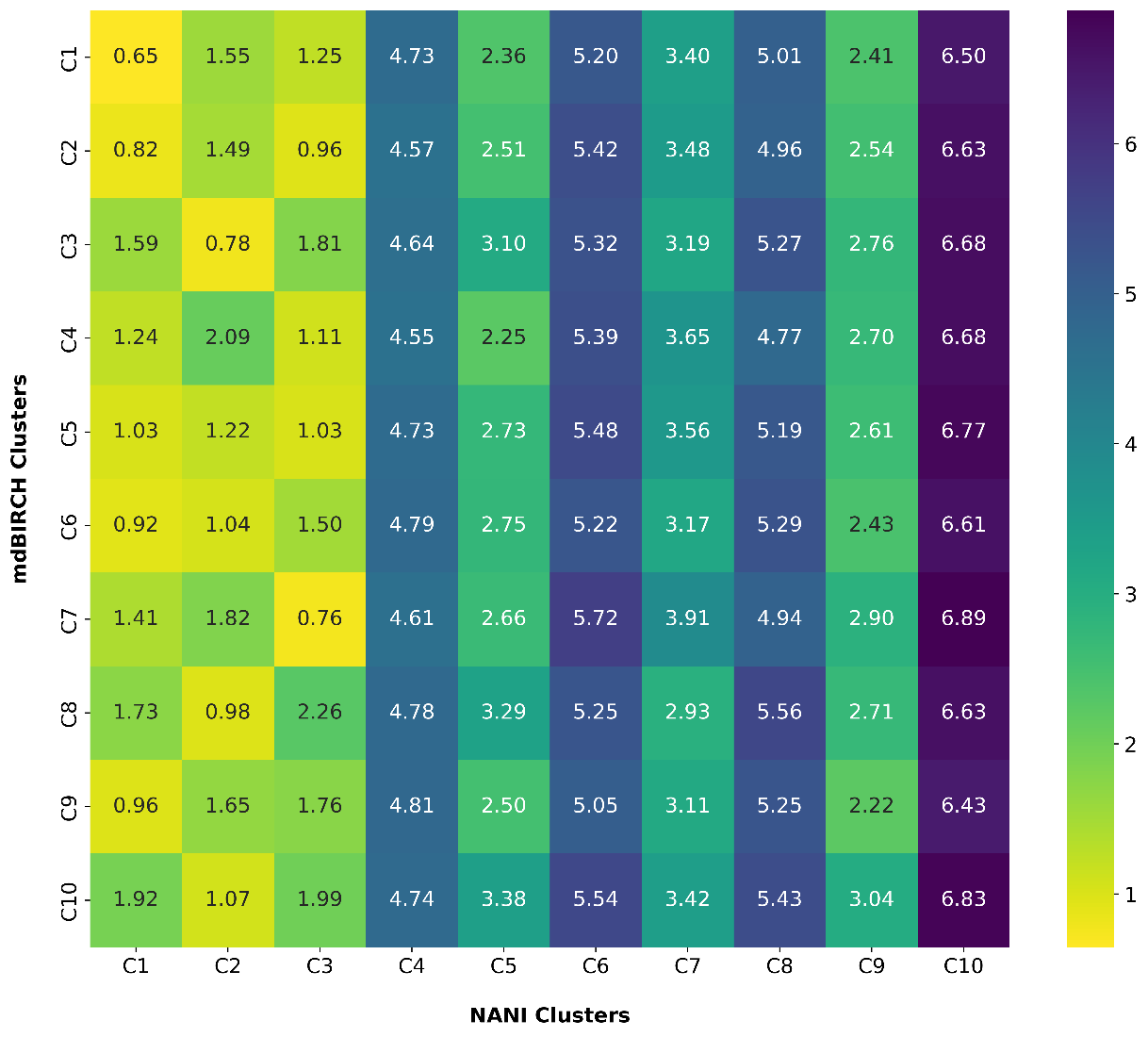

**Figure S5.** Medoid-to-medoid RMSD matrix comparison on the HP35 system. The heatmap reports pairwise RMSDs (Å) between cluster medoids from mdBIRCH (rows; the 10 most populated clusters, BF=1000, $\tau$=4 Å) and *k*-means NANI (columns; $k$=10, strat_reduced initialization type). Lighter colors (yellow) indicate lower RMSD and better structural agreement.
